# Urban crops demonstrate comparable potential for organic carbon storage as natural ecosystems

**DOI:** 10.1101/2025.01.15.633129

**Authors:** Beatriz Jiménez-Prieto, Pablo García-Palacios, Carmen Lorenzo, Elena Aguilar-Santana, Camelia Algora, Cristina Armas, Felipe Bastida, Leonor Calvo, Nuria Casado Coy, Axel Campos-Castro, Giada Centenaro, Sonia Chamizo, Joana Costa, Santiago Martín-Bravo, Manuel Delgado-Baquerizo, Svetlana Dashevskaya, María Dolores Carmona-Yáñez, Jorge Durán Humia, Maria José Fernández-Alonso, Daniela Figueira, Eva Garcia, Josu G. Alday, Enrique G. de la Riva, Mónica Ladrón de Guevara, Ana López-Velasco, Manuel Esteban Lucas-Borja, Ivan Prieto Aguilar, Jesús Pérez-López, Pedro Antonio Plaza-Alvárez, Alexandra Rodríguez Pereiras, Carlos Sanz-Lazaro, Santiago Soliveres, Alejandro Terrones, Aurora Torres, María Leo

## Abstract

Urban greenspaces, encompassing parks, golf courses, roundabouts, and urban crops, have potential to offset urban carbon footprints by storing soil organic carbon (SOC). This study analyzed particulate organic carbon (POC) and mineral-associated organic carbon (MAOC) stocks in topsoil across 27 Iberian cities, comparing urban greenspaces with natural ecosystems under varying climatic and edaphic conditions. Results revealed that urban greenspaces store comparable SOC stocks to natural ecosystems, with MAOC stocks being more dominant and stable across land-use types. POC stocks showed variability, particularly lower in roundabouts and parks compared to natural ecosystems, but remained similar in golf courses and urban crops. SOC fractions correlated inversely with mean annual temperature (MAT), emphasizing the need for cooling strategies in urban areas to preserve stable carbon pools. While MAOC exhibited saturation at higher SOC levels, POC showed a linear increase. This study highlights the importance of tailored management practices to enhance carbon storage in urban soils for climate change mitigation.

## Introduction

Soils represent the largest terrestrial reservoir of organic carbon (Lal, 2024). Preserving and increasing soil organic carbon (SOC) stocks is a promising natural climate solution to mitigate anthropogenic climate change (IPCC, 2021 ; Fuss et al., 2020). While SOC stocks will be eventually released to the atmosphere by microbial decomposition (Lal, 2024), SOC storage can help us buy time till anthropogenic emissions are dramatically reduced (Fuss et al., 2020). Increasing SOC stocks for climate change mitigation has been traditionally addressed in natural ecosystems such as forests, grasslands, and wetlands (Minasny et al., 2017; Paustian et al., 2016), and in human-altered ecosystems such as agricultural fields (Lal, 2024). However, the potential of urban greenspaces to store large amounts of SOC is understudied, despite the increasing importance of these areas in a context where 68% of the global population will live in cities by 2050 (United Nations, 2019). Comparing the ability of urban greenspaces to store SOC with that of their natural counterparts, as well as addressing the main climate, soil, and management drivers, is capital to manage the potential of these ecosystems to offset the C footprint of urban areas.

Despite spatially explicit models and meta-analysis suggest a direct impact on C pools with urban expansion (Seto et al., 2012; Chiens and Krumins 2022 STOTEN), recent global estimates based on a network of field sites found similar contents of SOC in urban greenspaces and nearby natural ecosystems (Delgado-Baquerizo et al 2022 Nat Clim Change). However, urban greenspaces host a diverse range of land use types with different management practices and intrinsic features that may alter their capacity to store SOC. For instance, golf courses are an example of intensively managed areas, often receiving inorganic fertilizers and high irrigation rates that might increase soil carbon inputs but also lead to SOC depletion due to constant mowing and compacted soils (Lal, 2004). In contrast, parks, with more natural vegetation and typically lower fertilization rates, might store higher levels of SOC due to the presence of a mix of herbaceous plants and woody species, with less intense human intervention (Minasny et al., 2017).

Despite most large-spatial scale assessments of soil C have focused on overall SOC stocks (refs), SOC is not in one piece, and may be conceptually categorized into particulate (POC) and mineral-associated (MAOC) organic carbon (Six et al., 2002). POC is the most susceptible fraction to climate and land use changes (ref), since it is not occluded in micropores or microaggregates and/or bound to mineral surfaces that protect organic C from microbial decomposition, as it is the case in the MAOC fraction (Cotrufo et al., 2019 + 1-2 more refs). The relative dominance of one soil fraction over the other may determine the capacity of urban greenspaces to accumulate C, since the C saturation hypothesis predicts a limit when adsorption sites on soil particles, such as clays, iron oxides, and aluminum oxides become filled, thereby limiting the soil’s ability to store additional carbon in the MAOC fraction (Kirkby et al., 2011; Rumpel & Kögel-Knabner, 2011). Disentangling the physical distribution of overall SOC stocks in the POC and MAOC fractions across a gradient of land use types and environmental conditions can inform management towards soil C stewardship, as performed in other terrestrial ecosystems (Cotrufo et al. 2019, García-Palacios et al. 2024, etc).

As the significance of urban greenspaces for climate change mitigation becomes increasingly recognized, it is essential to understand the main climatic features, soil parameters and management practices that determine SOC stock in these areas. Significant differences in carbon storage can be explained by the climatic characteristics and vegetation of each ecosystem (Jiang et al., 2017). Climatic conditions, such as increases in temperature and precipitation, can enhance soil organic carbon (SOC) by favoring plant productivity and biomass (Jiang et al., 2017; Chen et al., 2018). This is due to the fact that climate largely determines vegetation productivity and cover, and the amount of SOC is controlled by aboveground biomass and plant composition (Fornara and Tilman, 2008; Steinbeiss et al., 2008). This suggests that mitigation strategies based on C sequestration should not only focus on natural environments but also consider the potential of urban green spaces, as carbon storage associated with various urban land uses (such as golf courses, roundabouts, urban crops, and parks) remains largely unknown.

In this study, we addressed the stocks of POC and MAOC in the topsoil (10 cm) of four urban greenspaces (golf courses, roundabouts, urban crops, and parks) and one natural ecosystem across 27 cities in the Iberian Peninsula. Our site selection encompassed a wide edaphoclimatic gradient, with soil ph x to x, soil sand content x to x, MAT x to x and MAP x to x. First, we compared POC and MAOC stocks in the four urban greenspaces with the reference natural ecosystem. Next, we assessed the potential environmental and management controls driving POC and MAOC stocks. Lastly, we explored the relationship between SOC and the POC and MAOC contents to determine whether any of these fractions exhibit saturation.

## Materials and Methods

### Study sites and experimental design

We sampled four urban land uses (urban crop, park, roundabout, and golf course) and one natural ecosystem in 27 cities throughout the Iberian Peninsula, totaling 133 sites (Fig. 1). The golf course in Barcelona, Zaragoza y Segovia x and x in x could not be sampled due to lack of permits These cities were selected to capture a broad gradient in climate and soil parameters (Table S1), such as with mean annual temperature (MAT, x to x °C), mean annual precipitation (MAP, x to x mm), soil pH (x to x), soil sand content (x to x 5). The population of all cities was at least 10,000 inhabitants, and the five land use types were located at less than 20 km from each other to foster consistent climatic conditions. The four urban greenspaces encompass different management intensities, from parks with moderate management, urban crops with sustainable farming practices, to golf courses and roundabouts that require frequent irrigation and maintenance. On the other hand, the natural ecosystem represents a reference for comparison with urban greenspaces, keeping climatic conditions constant.

**Figure 1.**
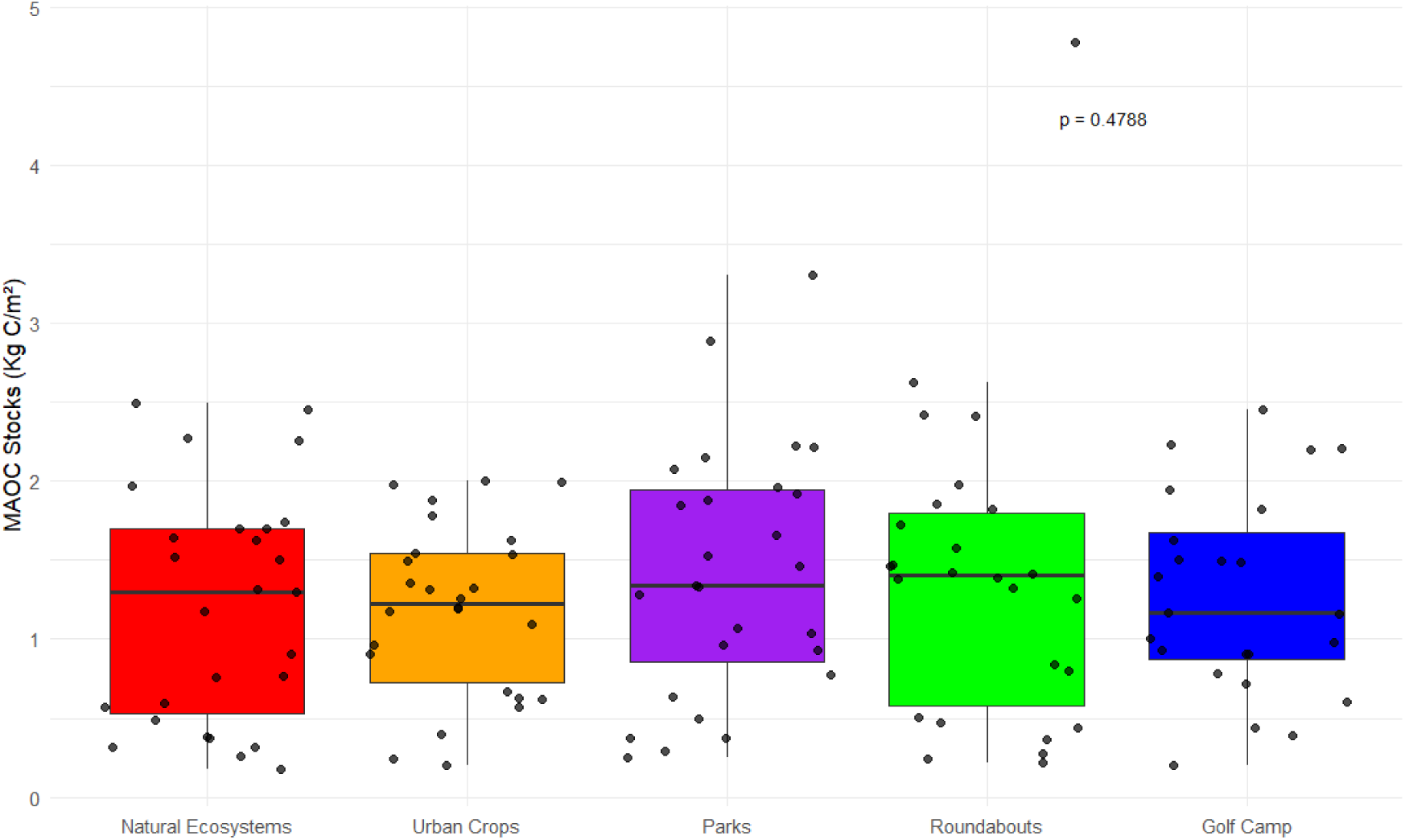
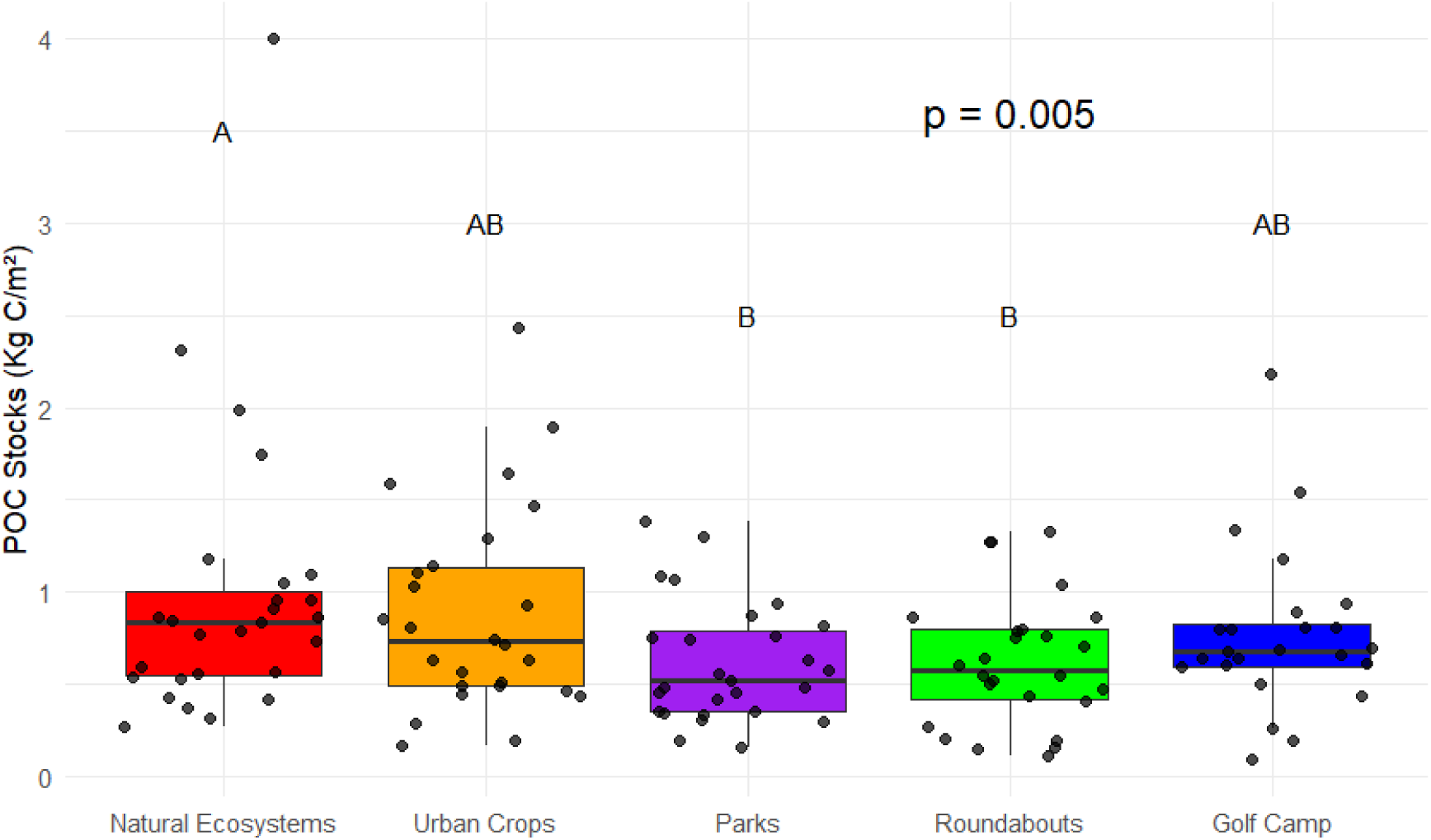
Soil POC (a) and MAOC (b) stocks in natural ecosystems compared with four urban greenspaces. NE (natural ecosystem, n = x), UC (urban crop, n = x), UG (urban garden, n = x), R (roundabout, n = x), and GC (golf course, n = x). Box plots represent first and third quartiles (box), medians (central horizontal line), largest value smaller than 1.5 times the interquartile range (upper vertical line) and smallest value larger than 1.5 times. Letters represent significant differences at *P* < 0.05 in post-hoc analysis.

### Field sampling

We established a 20× 20 m plot at each site, with three 12 m-long transects per plot and three 4 × 4 m subplots per transect. To account for spatial heterogeneity, a composite surface soil sample (top 10 depth) was collected using nine soil cores from the subplots. We focused on surface soils because it is the most biologically active layer (Rey, Pegoraro, & Jarvis, 2008), it is particularly important in a context of climate and land use change (Crowther et al., 2016; Jobbágy & Jackson, 2000), and it allows comparison with previous soil C studies across large environmental gradients (Cotrufo et al., 2019, Delgado-Baquerizo, García-Palacios, Bradford, et al., 2023). Also, urban greenspaces typically have shallow soils as a consequence of surface preparation and maintenance (ref). Soil samples were sieved at 2 mm and roots were removed when present. An additional 10-cm depth soil core was collected to determine bulk density (g cm^-3^) as the mass of dry, coarse fragmented-free soil per volume of excavated soil. We recorded total plant cover and plant diversity (Shannon index using the relative abundance of x plant functional groups such as herbs, shrubs, trees, etc) as surrogates of management determining plant C inputs quantity and quality, respectively. We also collected climate information from WorldClim 2.2 (Fick & Hijmans, 2017) using spatial coordinates from each site, specifically MAT and MAP. This climatic database, which covers the period from 1970 to 2000, also incorporates more recent high-resolution data derived from a wide array of climatic observations and topographic data from the Shuttle Radar Topography Mission (SRTM).

### Laboratory analyses

To determine the stocks of soil organic C stored as MAOC and POC, a physical fractionation procedure was conducted using a 53 µm sieve. A 10.5 g sample of each soil was agitated in a sodium hexametaphosphate solution (5 g/L) for 18 hours to ensure complete dispersion. Subsequently, an automatic fractionator was employed to separate the two soil fractions by passing distilled water through the 53 µm sieve. The fraction that passed through was collected as mineral-associated organic matter, while the fraction retained on the sieve was referred to as particulate organic matter. After drying the fractions to a constant weight in an oven at 60 °C, the total content (%) of organic C in each fraction and in bulk soil samples was measured using an Elemental Analyzer (C/N Flash EA 112 Series-Leco Truspec). Carbonates were removed by acid fumigation before analysis. Using SOC, POC and MAOC contents, soil bulk density and soil core depth, we calculated SOC, POC and MAOC stocks in kg m^-2^. To measure soil pH, a suspension was prepared in a 1:10 (mass) ratio of soil to water, and a calibrated pH meter was used for accurate measurement.We selected the percentage of sand for subsequent analysis because it is considered the most adequate variable to address the relative influence of soil texture on C storage and ecosystem functioning (Maestre et al., 2012; García-Palacios et al. 2024 Nat Geo).

### Data analyses

We used linear mixed-effects modelling to compare SOC, POC and MAOC stocks (kg m^-2^) in natural ecosystems with urban land use types (urban crop, park, roundabout, and golf course). At each of the three models, land use type (five levels) was included as a fixed factor, while city was included as a random factor to capture unexplained variability between cities. A post-hoc analysis was conducted to evaluate whether the differences among levels of land use type were significant. To investigate the relative influence of climate, soil parameters, and management on POC and MAOC stocks (kg m^-^2) across the four urban ecosystems, two linear mixed models were performed, one per fraction type. In these models, climate variables (MAT and MAP), soil parameters (pH and sand content), and management factors (tree cover and plant diversity) were included as fixed factors, while city was included as a random effect. To model the non-linear relationship between POC and MAOC contents (%) with SOC, two generalized additive models (GAM) were conducted, one per fraction type. All the analyses were conducted using R version 4.4.1 and the lme4 (Bates et al. 2015), emmeans (Lenth, 2021), and mgcv (Wood, 2017) packages.

## Results

### Differences in POC and MAOC stocks in natural ecosystems and urban greenspaces

POC stocks changed with land use type (p = 0.0050, Fig. 1a). The post-hoc analysis revealed that natural ecosystems stored x% (median: x kg m^-2^; interquartile range: x) and x% (median: x kg m^-2^; interquartile range: x) more POC stocks than roundabouts and urban gardens, respectively, but stocks were similar than in golf courses and urban crops. In contrast, MAOC stocks were similar across the five land use types (p = 0.4788, Fig. 1b). The mean MAOC stocks across land use types (1.298 kg m^-2^) were 68% higher than POC stocks (0.775 kg m^-2^). Overall SOC stocks, measured in bulk soil samples, did not change with land use type (p = x, Fig. S1).

### Environmental and management drivers of POC and MAOC stocks across urban greenspaces

The variance explained by the fixed-effects predictors was x% and x% for POC and MAOC stocks, respectively (Fig. 2). Overall, the climatic, soil and management factors evaluated did not influence POC and MAOC stocks. We only found a significant decrease in both POC (estimate: x, 95% CI: x to x) and MAOC (estimate: x, 95% CI: x to x) stocks with MAT. Higher POC stocks were associated with lower soil pH (estimate: x, 95% CI: x to x), plant diversity (estimate: x, 95% CI: x to x) and MAP (estimate: x, 95% CI: x to x), although these effects were not significant.

**Figure 2.**
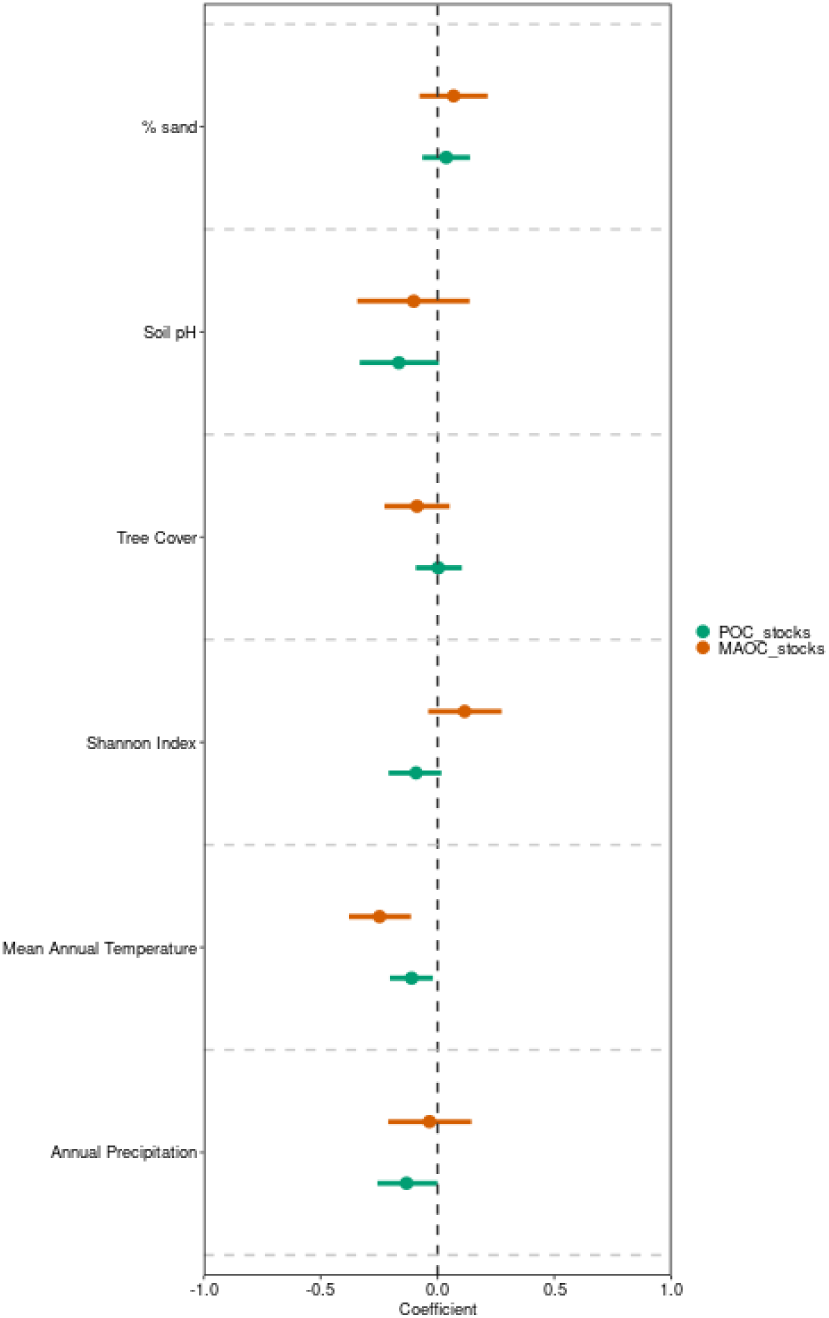
Effects of climate, soil parameters and management on soil POC and MAOC stocks across four urban greenspaces (urban crop, urban garden, roundabout and golf course). The dots and lines represent the coefficients and 95% Cis of the fixed effects of the predictors evaluated in the two linear-mixed effects models (n POC model = x, n MAOC model = x).

### The relationships between SOC and contents of POC and MAOC in natural ecosystems and urban greenspaces

When addressing this relationship across the four urban greenspaces, both fractions exhibited a non-linear relationship with overall SOC content (Fig. 3). However, the shape of this relationship was different in POC than in MAOC. While POC content steadily increased with SOC (P = x, Fig. 3a), the increase in MAOC content saturated at higher SOC values (P = x, Fig. 3b). As a consequence, MAOC dominance relative to POC decreased at higher SOC. When individual urban greenspaces were compared with natural ecosystems, we found that the saturating relationship between MAOC and SOC was particularly evident in natural ecosystems, parks and golf courses (all P < 0.05, Fig. 4a), while POC did not saturate in any of the land use types (all P < 0.05, Fig. 4b).

**Figure 3.**
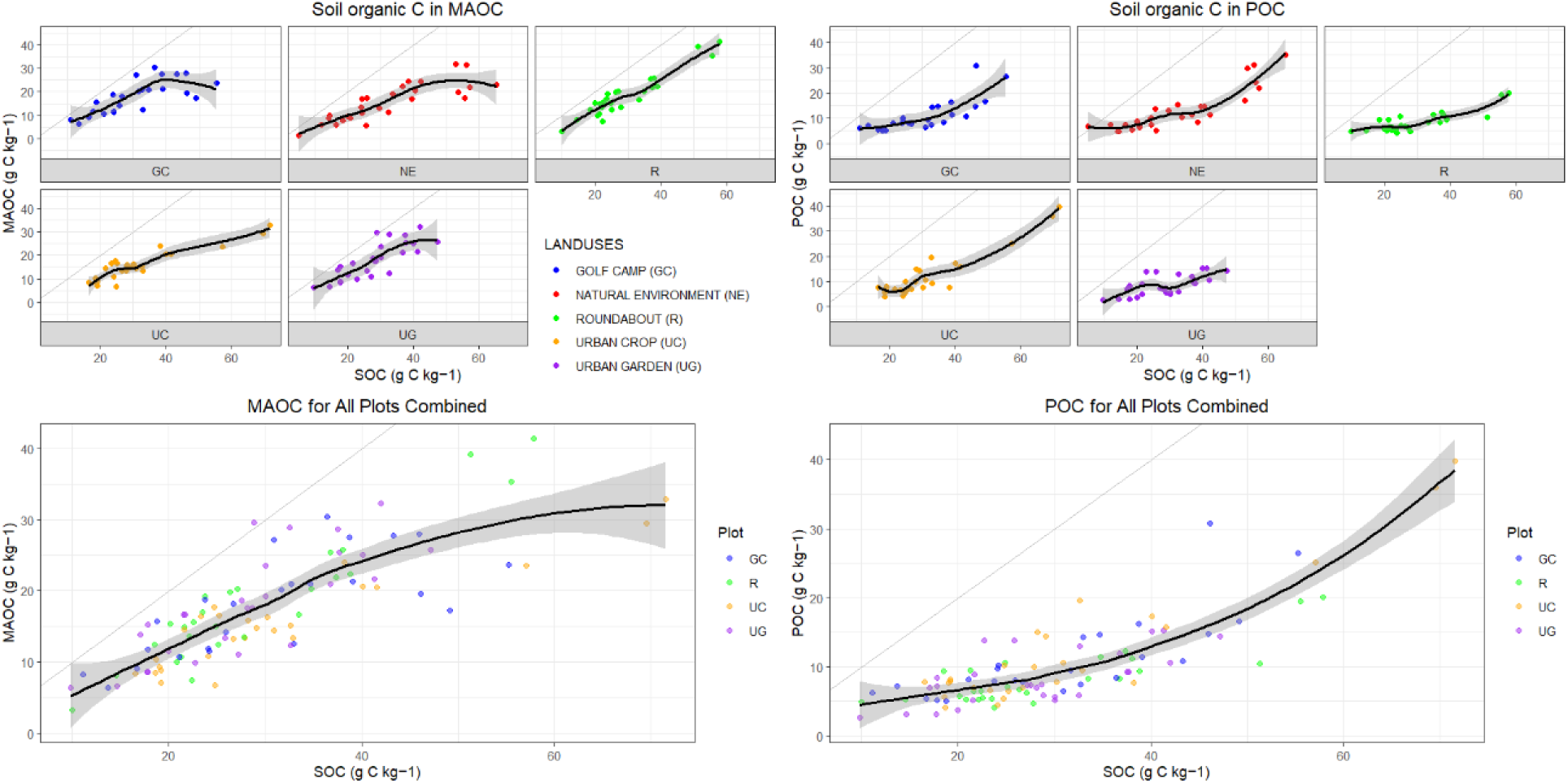
Relationships between SOC and contents of POC (a) and MAOC (b) across four urban greenspaces (urban crop, urban garden, roundabout and golf course). Lines and shadings represent the non-linear relationships in generalized additive models (GAM) and 95% CI. n = x and x in the POC and MAOC models.

**Figure 4.**
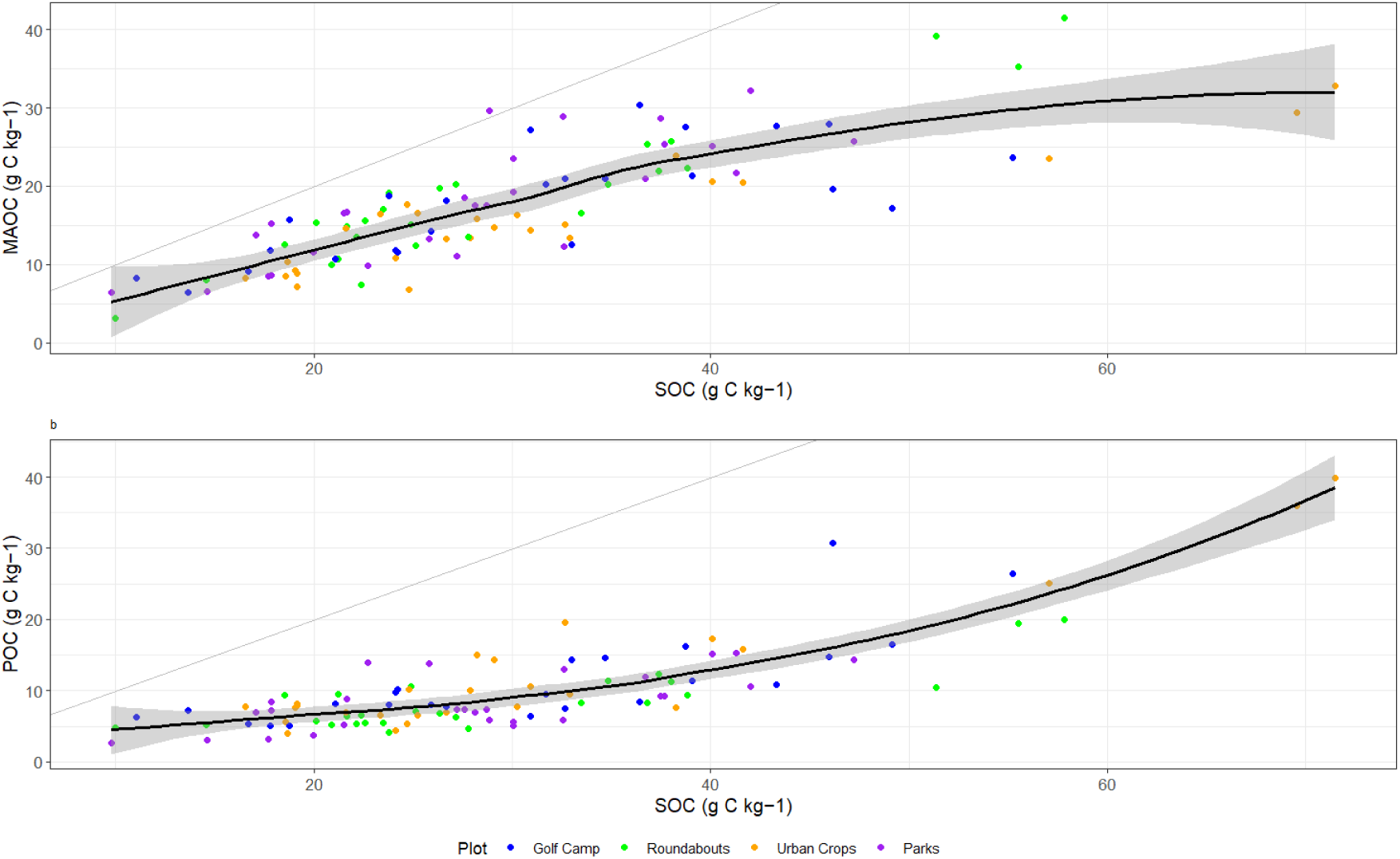
Relationships between SOC and contents of POC (a) and MAOC (b) in natural ecosystems compared with four urban greenspaces (urban crop, urban garden, roundabout and golf course). Lines and shadings represent the non-linear relationships in generalized additive models (GAM) and 95% CI. n = x, x, x, x and x (NE, UG, UC, R and GC) in the POC models and n = x, x, x, x and x (NE, UG, UC, R and GC) in the MAOC models.

## Discussion

The physical fractionation of soil organic matter has been proposed as a meaningful framework to inform land management towards climate change mitigation (Cotrufo et al., 2019). Urban greenspaces can help to offset the C footprint of urban areas if substantial C can be stored in their soils for a relevant amount of time. Our observational study across 27 cities spanning a wide soil-climatic gradient in the Iberian Peninsula found that urban greenspaces stored a similar stock of MAOC, the less sensitive SOC fraction to climate and land use changes, than natural ecosystems. Also, POC stocks in golf courses and urban crops paralleled POC stocks in natural ecosystems. POC and MAOC stocks were consistently higher in colder sites, but other key drivers of soil C such as precipitation, soil pH, soil sand, plant diversity and cover were poorly related with both C fractions. These findings underscore the potential of urban greenspaces to store large amounts of soil C in the POC and MAOC fractions.

The four urban greenspaces considered, golf courses, urban crops, parks and roundabouts, stored a similar amount of SOC per unit area than nearby natural ecosystems, confirming the results found by Delgado-Baquerizo et al. (2023) at the global scale, and suggesting that urbanization has not largely altered the land C sink. Importantly, the absence of SOC differences between natural and urban greenspaces was driven by the lack of response of MAOC to changes in land use type, and the overall dominance of MAOC over POC stocks.

Although MAOC stocks substantially varied across the environmental gradient assessed, ranging from x to x kg C m2 (Fig. 1a), these changes were not related with the identity of the land use type. Soil C stored in the MAOC fraction is protected from microbial decomposition because organo-mineral associations and occlusion within aggregates limit the accessibility of microorganisms and their extracellular enzymes to the organic substrates. Our study extends previous findings on the lower sensitivity of MAOC compared to POC to climate change and agricultural conversion, to the urbanization process. These results suggest that the MAOC fraction is less affected by urban soil management practices such as irrigation, compaction, or organic residue management (Stewart et al., 2007). Furthermore, MAOC were 68% higher than POC stocks across the five land use types evaluated, which is in line with the MAOC dominance found in temperate and dryland biomes that characterize our sampled sites in the Iberian Peninsula (Sokol et al 2022 Functional Ecology, Díaz-Martínez et al. 2024 Nature Clim Change).

In contrast, POC stocks were significantly lower in roundabouts and parks than in natural ecosystems, but remained the same in golf courses and urban crops (Fig. 1b). Although the four urban greenspaces are intensively managed, with constant irrigation and mowing, differences in practice may drive this pattern. We speculate that high annual rates of organic amendments in golf courses and urban crops via mulching and manure (García-Orenes et al., 2012) may help to maintain POC levels as high as in natural ecosystems. The addition of fresh organic inputs and the cascading effects reducing soil compaction for better aeration and microbial activity are both crucial for POC formation (Díaz et al., 2020, López-Ballesteros et al., 2020). Taken together, our findings suggest that urbanization has not altered the capacity of soil ecosystems to store large amounts of SOC, particularly in the MAOC fraction.

Temperature is one of the most prominent controls driving the spatial distribution of SOC in natural ecosystems, with a very clear pattern towards higher content in colder regions (García-Palacios et al. 2021 NREE). Our results show that POC and MAOC stocks decrease with MAT in the four urban greenspaces evaluated. Although the geographic coverage of the Iberian Peninsula is reduced, it represents a wide MAT gradient, from X to X °C. This result is in line with major findings from global biomes (Conant et al., 2011, Diaz-Martínez et al 2024 Nat Clim Change), but most importantly, agrees with the results of a global network of urban ecosystems (Delgado-Baquerizo et al. 2023). In particular, the negative response of MAOC to temperature highlights the importance of implementing warming mitigation strategies in urban areas to preserve the more persistent C reserves in soil. Despite its stability, MAOC can still be affected by climate change, influencing its release and storage in the soil (Lehmann & Kleber, 2015). Other studies have shown that urban soils can have surprisingly high capacities for long-term C storage, especially when appropriate management strategies are applied (Edmondson et al., 2014). If lower MAT drive the accumulation of POC and MAOC in urban soils, warming these areas is likely to reverse this effect, and mineral binding and occlusion within aggregates may not counteract such C losses.

Beyond MAT, the rest of the climatic, soil and management features evaluated were poorly related with POC and MAOC stocks, despite the wide gradients in MAP (x to x mm), soil pH (x to x), soil sand (x to x) and plant cover (x to x). This was particularly remarkable for MAOC, especially regarding its lack of sensitivity to variation in sand content, as MAOC usually accumulates in clayey soils with large mineral surface area to interact with. Also, POC and MAOC stocks seem to be decoupled from increases in plant C inputs as addressed by tree cover, which is a major SOC stabilization mechanisms in natural ecosystems (Cotrufo et al., 2013). POC was somehow influenced by lower plant diversity and MAP. This counterintuitive result may be driven by the higher POC content found in golf courses and urban crops, which are inherently dominated by a few plant species.

Theory predicts that the accumulation of C in the POC fraction steadily increases as long as C inputs are not limiting, while MAOC is dependent on a finite availability of mineral surfaces to interact with (Stewart et al. 2008 SSAJ, Cotrufo et al. 2023 GCB). This may explain the saturating MAOC response found in natural ecosystems (Cotrufo et al. 2019 Nat geo, García-Palacios et al. 2024 Nat Geo), although recent studies have challenged this assumption (Begill et al. 2023 GCB). Here we explored the linkages between overall SOC and POC and MAOC fractions with GAM models, to capture non-linear relationships, and found evidence for such a MAOC saturating response in urban greenspaces. For instance, POC content across urban greenspaces monotonically increased with SOC, but MAOC increases levelled off at higher SOC contents. Thus, MAOC dominance over POC decreases at SOC higher than 50 g C kg^-1^, although the sample size is low at this side of the gradient, considerable increasing the error associated with our estimates. Individually, natural ecosystems, parks and golf courses exhibited a saturating MAOC relationship, suggesting that this response is particularly evident in the land use types with larger SOC content, where there is more potential to reach the availability of mineral surfaces (Cotrufo et al. 2019 Nat geo, García-Palacios et al. 2024 Nat Geo).

## Conclusion

In summary, we evaluated the capacity of various urban green spaces (golf courses, roundabouts, urban gardens, and parks) to store carbon in 27 cities across the Iberian Peninsula. Our results revealed a variable potential for carbon storage, influenced by land use and temperature. Although urban green spaces tend to store less particulate organic carbon (POC) than natural ecosystems, urban gardens exhibit POC reserves comparable to those of natural ecosystems. This study demonstrates that urban green spaces contain significant surface carbon stocks across the edaphoclimatic gradient of the Iberian Peninsula. While previous research has focused on carbon in natural ecosystems (Minasny et al., 2017; Paustian et al., 2016), these results represent a significant advance in our understanding of carbon storage in urban ecosystems. Our analysis of particulate organic carbon (POC) and mineral-associated organic carbon (MAOC) further clarifies the mechanisms and limitations of carbon storage in these spaces. By breaking down the specific dynamics of carbon, as well as its POC and MAOC fractions, in different urban green spaces, this study establishes a foundation for future research. Our findings highlight the importance of implementing soil organic carbon (SOC) management strategies tailored to each type of urban green space, based on local edaphoclimatic factors. The results also demonstrate the negative effect of temperature on MAOC stocks. This suggests that proper planning and management of urban spaces in the context of climate change could optimize carbon sequestration in these areas. Despite the increasing importance of urban green spaces in the context of global urbanization, their potential for SOC storage has been underexplored. Given the ongoing urban expansion, maximizing the potential of urban green spaces as carbon reservoirs is a key strategy for mitigating climate change in urban areas, providing a basis for developing more sustainable urban management models and policies that maximize environmental benefits in cities.

## Notes

### Competing Interest Statement

The authors have declared no competing interest.

